# Epithelial Yap/Taz are required for functional alveolar regeneration following acute lung injury

**DOI:** 10.1101/2023.06.22.545997

**Authors:** Gianluca T. DiGiovanni, Wei Han, Taylor Sherrill, Chase J. Taylor, David S. Nichols, Natalie M. Geis, Ujjal K. Singha, Carla L. Calvi, A. Scott McCall, Molly M. Dixon, Yang Lui, Ji-Hoon Jang, Sergey S. Gutor, Vasiliy V. Polosukhin, Timothy S. Blackwell, Jonathan A. Kropski, Jason J. Gokey

## Abstract

A hallmark of idiopathic pulmonary fibrosis (IPF) and other interstitial lung diseases is dysregulated repair of the alveolar epithelium. The Hippo pathway effector transcription factors YAP and TAZ have been implicated as essential for type 1 and type 2 alveolar epithelial cell (AT1 and AT2) differentiation in the developing lung, yet aberrant activation of YAP/TAZ is a prominent feature of the dysregulated alveolar epithelium in IPF. In these studies, we sought to define the functional role of YAP/TAZ activity during alveolar regeneration. We demonstrate that Yap and Taz are normally activated in AT2 cells shortly after injury, and deletion of Yap/Taz in AT2 cells led to pathologic alveolar remodeling, failure of AT2 to AT1 cell differentiation, increased collagen deposition, exaggerated neutrophilic inflammation, and increased mortality following injury induced by a single dose of bleomycin. Loss of Yap/Taz activity prior to a LPS injury prevented AT1 cell regeneration, led to intra-alveolar collagen deposition, and resulted in persistent innate inflammation. Together these findings establish that AT2 cell Yap/Taz activity is essential for functional alveolar epithelial repair and prevention of fibrotic remodeling.

## Introduction

Since the lung is subjected to a large number of injurious agents throughout life including viral infection, tobacco smoke, and environmental particulates, the capacity for functional repair is essential for maintenance of normal gas exchange. The alveolus, which is responsible for gas exchange, is comprised of two primary types of alveolar epithelial cell. Alveolar type 1 (AT1) cells are large, thin, irregularly shaped cells that normally are located in close proximity to alveolar capillaries and facilitate gas exchange in the lung. The alveolar type 2 (AT2) cell secretes pulmonary surfactant and functions as a facultative progenitor cell for AT1 cells, thus supporting both alveolar structure and function ^1, 2^. Injury to AT1 and AT2 cells requires proliferation and differentiation of progenitor cells to reestablish the epithelial barrier and regenerate functional alveoli. The mechanisms directing this initial proliferative response followed by differentiation into functional AT1 cells remain incompletely understood. Recent work from our group and others has implicated the Hippo-Yap/Taz signaling pathway as a regulator of alveolar epithelial development and repair ^3–5^, but also indicated that dysregulation of this pathway is a prominent feature of the distal lung epithelium in idiopathic pulmonary fibrosis ^6^.

The Hippo-Yap/Taz core pathway consists of components mammalian serine threonine kinases *MST1/2* (referred to as serine threonine kinases *Stk4/3* respectively in the mouse), that phosphorylate large tumor suppressor kinases (LATS1/2), that then phosphorylates transcriptional co-factors Yap and Taz (*Wwtr1*). In the presence of this phosphorylation cascade, Yap and Taz are sequestered in the cytoplasm where they can undergo 14-3-3 ubiquitination and degradation. In the absence of the Hippo phosphorylation cascade, Yap and Taz shuttle to the nucleus where they interact with binding partners TEADs 1-4 ^7, 8^, as well as Runx2 and Smad2/3, to activate transcription for target genes ^9, 10^.

In the airway epithelium, there is evidence that temporally regulated Yap activity is required to maintain homeostasis. Sustained activation of Yap leads to basal cell metaplasia, whereas Yap deletion accelerates terminal differentiation of basal cells and impairs basal cell progenitor function, indicating a tight regulation of Yap activity during airway development and repair ^11–14^. During alveologenesis, ectopic Yap/Taz activation leads to increased AT2 cell proliferation and increased number of AT1 cells (as defined by Hopx+/Ager+ expression), while Yap deletion results in decreased proliferating AT2 cells and decreased number of AT1 cells ^4, 5^. We and others have shown that Yap deletion in AT2 cells increases “mature” AT2 cell marker expression, including *Sftpc* and *Abca3* ^4, 15, 16^. These findings collectively provide evidence that Yap/Taz activity regulates alveolar epithelial cell fate specification during postnatal lung development. However, there has been limited investigation of the Hippo-Yap/Taz pathway during alveolar repair in the adult lung ^3^.

Herein, we sought to understand the dynamics of Yap/Taz activity during normal alveolar repair and establish whether Yap/Taz activity is required for functional alveolar regeneration following injury. Using a genetic model of AT2 cell specific deletion of Yap and Taz, mouse lungs were injured with intratracheal delivery of bleomycin or lipopolysaccharide (LPS). We found that Yap/Taz deletion leads to persistent alveolar inflammation, and results in exaggerated fibroblast activation and lung fibrosis in both bleomycin and LPS injury models. These findings demonstrate that AT2 cell specific Yap/Taz deletion is essential for initiation of alveolar repair and without Yap/Taz activation there is a failure to regenerate AT1 cells necessary for functional alveoli.

## Results

### Alveolar epithelial Yap/Taz activity exhibits cell-type specific dynamic changes during lung injury and repair

To understand the dynamics of Yap/Taz activity during alveolar injury and repair, wild-type C57Bl6 mice were challenged with a single dose of intratracheal (IT) bleomycin (0.08 IU) and sacrificed 4, 7, 14 and 21 days after bleomycin injury (**Figure 1A).** In control mice, nuclear Yap or Taz was rarely detected in Sp-C+ AT2 cells (**Figure 1 B, E, G**) and nuclear Yap was detected in a small minority of Hopx+ AT1 cells (5.1 ±1.4%); however, nuclear Taz was present in nearly all (97.9± 1.1%) AT1 cells (**Figure 1 F, H**). After bleomycin-induced injury, quantification of Sp-C+ AT2 and Hopx+ AT1 cells revealed both cell types are reduced by day 4 and return to baseline numbers by 21 days post-injury (**Figure 1 C,D**). At 7 days after bleomycin, nuclear Yap and Taz were found in 72.7 ± 2.9% and 12 ± 2.1% of Sp-C+ cells respectively (**Figure 1**). At 14 days after injury, 40.0 ± 2.2% of Sp-C+ AT2 cells had nuclear Yap, which decreased further to 13.3 ± 1.5% by day 21. In comparison, nuclear Taz was identified in 35.2 ± 4.2% of AT2 cells at day 14 and to 23.1 ± 2.8% at 21 days post-injury. Yap was present in approximately 10-13% of Hopx+ AT1 cells during repair (days 4, 7, and 14) and returned to 6.7 ± 1.6% by day 21. In contrast, Taz was readily detected in >93% of Hopx+ AT1 cells throughout homeostasis and repair, with the exception of 14 days post-injury where Taz was detected in 86.2 ± 1.9% of Hopx+ AT1 cells. Quantitative qPCR analysis of Yap/Taz targets in sorted Cd326+ epithelial cells identified a similar trend in RNA levels of *Ctgf, Axl,* and *Ajuba,* as targets are highest expressed by day 7 after injury and return to baseline levels by day 21 **(Figure 1I-K)**. These findings indicate that alveolar injury is followed by an asynchronous activation of Yap/Taz in regenerating alveolar epithelial cells peaking at days 7 and 14 respectively, followed by downregulation of Yap and stabilization of AT1 cell Taz as the lung returns to homeostasis.

**Figure 1:**
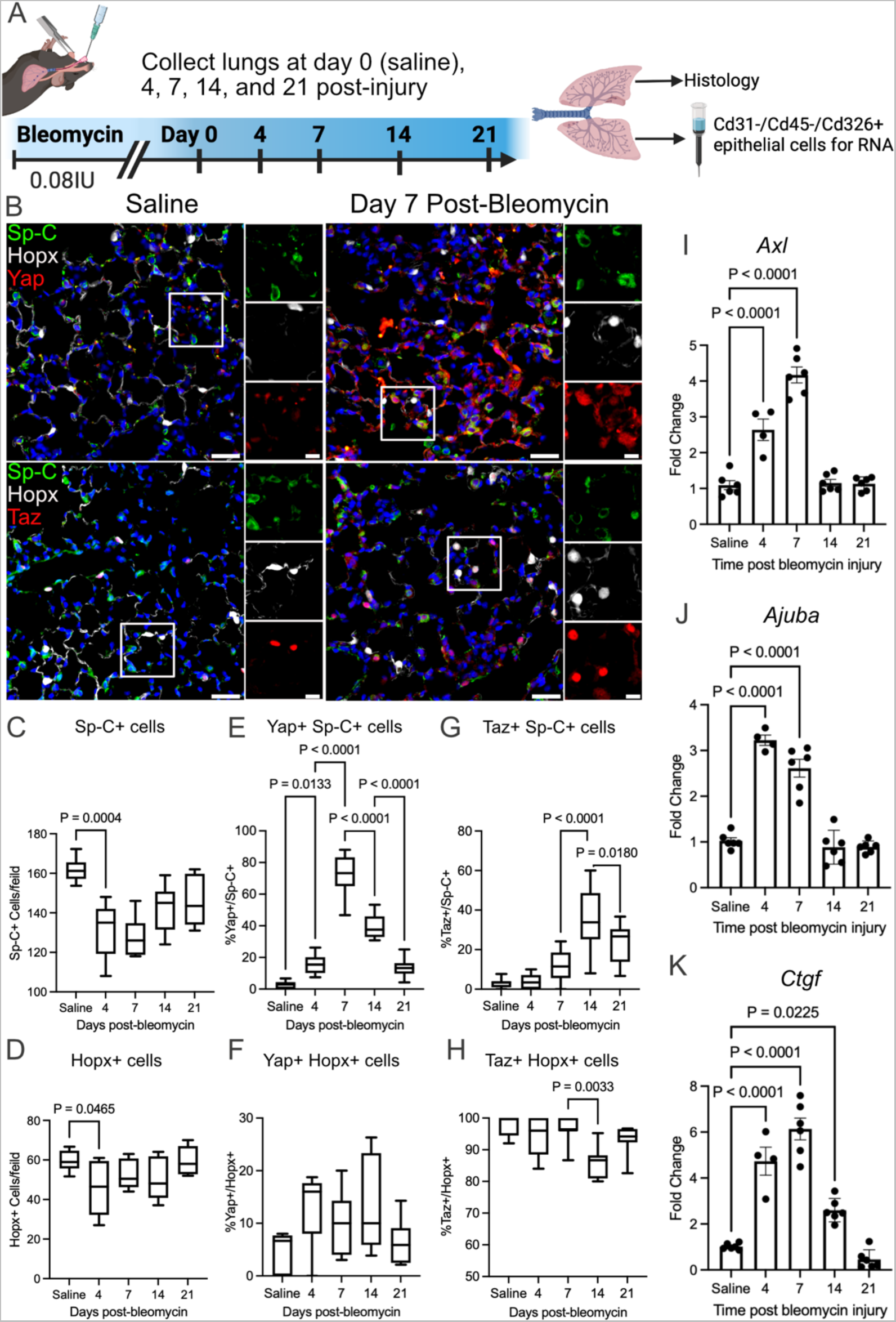
YAP/TAZ are dynamically regulated during acute bleomycin lung injury. **A)** Schematic of injury model and time-points when lungs were assessed. **B)** Immunofluorescence analysis of Yap or Taz (red) in Sp-C+ (green) AT2 and Hopx+ (white) AT1 cells. **C,D)** Quantification of Sp-C+ AT2 and Hopx+ AT1 cells at respective injury repair time-points. **E, F)** Quantification of Yap+ nuclei in Sp-C+ AT2 and Hopx+ AT1 cells. **G,H)** Quantification of Taz+ Sp-C+ AT2 and Hopx+ AT1 cells. **I,J,K)** qPCR analysis of Yap/Taz target genes **(I)** *Axl,* **(J)** *Ajuba,* and **(K)** *Ctgf* during bleomycin injury repair. Scale bars of low magnification images are 50μm and insets are 10μm. In box-whisker plots, whiskers are min and max and data represents N=6 mice, 10 frames imaged per mouse. Bar graphs show individual values with SEM error bars. Statistical analysis was performed using one-way ANOVA and Tukey’s post-hoc comparison.

*Yap is required for initiation of alveolar repair following bleomycin injury*.

Having observed early activation of Yap/Taz in AT2 cells following injury, we then sought to determine the role of AT2 cell Yap/Taz in regulating alveolar repair. Yap and Taz floxed mice (*Yap^flox/flox^Taz^flox/flox^*) were crossed with the AT2-cell specific promoter SftpcCreert2 ^17^ to drive tamoxifen inducible deletion of Yap and Taz (hereafter referred to as YT^del^ mice) in 8-10 week old adult mice 2-weeks prior to bleomycin induced lung injury (**Supplemental Figure S1A**). YT^del^ mice injured with IT bleomycin resulted in high mortality during the first 2 weeks post-injury. IT delivery of 0.08 IU/mouse resulted in approximately 75% mortality (**Supplemental Figure S1B**). Therefore, to establish a system in which the epithelial repair-repair response could be assessed, we treated mice with tamoxifen to delete Yap/Taz at the time of bleomycin injury (**Figure 2A**), which resulted in increased survival compared to the previous model with 50% mortality in YT^del^ mice compared to wild type Yap^flox/flox^Taz^flox/flox^ and C57/Bl6J control mice at 28 days post-injury (**Figure 2B**). Masson’s Trichrome staining (**Figure 2C**) and Sircol collagen analysis revealed that bleomycin-treated YT^del^ mouse lungs had increased soluble and total collagen compared to control mice (**Figure 2D**). YT^del^ mice had increased injured lung area as assessed by histological analysis of areas with collagen deposition or immune infiltrate ^18^, 28-days after bleomycin compared to WT sibling mice (**Figure 2E**). Immunofluorescence analysis showed that YT^del^ mice had reduced Hopx+ cells, while numbers of Sp-C+ cells remained similar to unchallenged mice 28-days after injury (**Figure 2F, G**).

**Figure 2:**
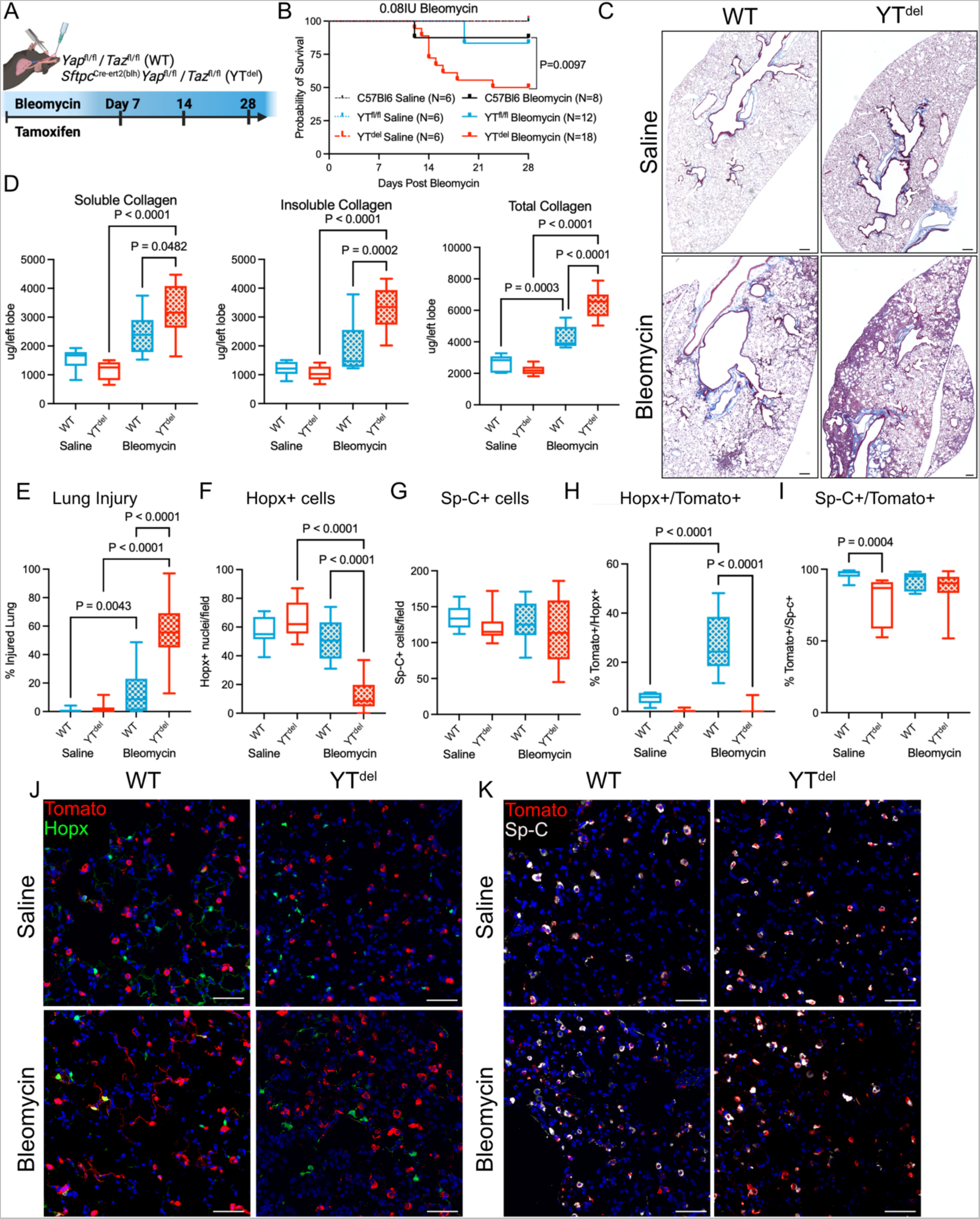
Deletion of YAP/TAZ leads to failed alveolar repair following single-dose bleomycin induced lung injury. **A)** Schematic of injury model in which mice were treated with tamoxifen the same day as bleomycin induced lung injury. **B)** Survival curve of wild-type and YT^del^ bleomycin or saline treated mice out to 28-days post-injury. Statistics determined using Mantel-Cox test. **C)** Masson’s Trichrome staining of tissue sections from WT and YT^del^ mice at day 28 after bleomycin or PBS. **D)** Soluble, insoluble, and total collagen quantification from WT and YT^del^ at day 28 after saline or bleomycin. **E)** Quantification of total injured lung area in respective groups at day 28 after saline/bleomycin. **F, G)** Quantification of total Hopx+ cells or Sp-C+ cells per 20X field of view in each group. **H, I)** Quantification of Sp-C^tomato^ lineage labeled Hopx+ or Sp-C+ AT2 cells 28-days after bleomycin. **J)** Immunofluorescence analysis of Sp-C^tomato^ lineage labeled cells (red) and Hopx+ (green) AT1 cells at 28-days post-injury. **K)** Immunofluorescence analysis of Sp-C^tomato^ lineage labeled cells (red) and Sp-C+ (white) AT2 cells at 28-days post-injury. Scale bars are (C) 200μm or (J,K) 50μm. Statistical analysis (D-I) was performed using one-way ANOVA and Tukey’s post-hoc comparison.

Observing reduced AT1 cell numbers in YT^del^ mice, we next sought to directly determine the ability of YT^del^ AT2 cells to act as progenitor cells to regenerate AT1 and AT2 cell populations. We performed lineage tracing studies using YT^del^ mice crossed with *Rosa26^lsl-tDtomato^*reporter mice, in which a Tomato fluorescent reporter is activated in AT2 cells when mice are treated with tamoxifen ^19^. While the number of lineage-labeled Sp-C cells (AT2 cells) was similar in bleomycin injured YT^del^ and WT mice, the number of lineage-labeled Hopx+ cells (AT1 cells) was reduced from 27.8 ± 3.3% in WT mice to 1.1 ± 0.4% in YT^del^ mice at 28-days post-injury (**Figure 2J, K).** Next, to test whether the effects of YT deletion on AT2 progenitor function were mediated through an autonomous mechanism, we isolated Cd45-/Cd31-/Cd326+ epithelial cells from lineage-labeled naïve WT and YT^del^ mice and established feeder-free organoids using recently described methodlology^20^ (**Supplemental Figure S2A**). By day 14, quantification of Tomato+ organoids revealed a dramatic reduction in the number (57.8 ± 3.9 compared to 5.1 ± 0.70) and size (77.8 ± 1.4 compared to 45.8 ± 1.9 μm) of organoids generated from YT^del^ AT2 cells compared to those generated with WT AT2 cells (**Supplemental Figure 2B, C**). Together, these findings demonstrate that loss of Yap and Taz in AT2 cells prevents AT2 to AT1 differentiation and impairs AT2 progenitor function.

### AT2 cell specific deletion of Yap/Taz leads to increased intermediate alveolar epithelial cell, activation of fibroblasts and increased immune cell populations

To investigate the mechanisms through which AT2-cell deletion of YT alters epithelial cell transcriptional programs and worsens experimental lung fibrosis, we then performed single-cell RNA sequencing of lung single cell suspensions generated from YT^del^ and wild-type saline and bleomycin treated mouse lungs at 28 days post injury (**Figure 3A**). Analysis identified 26 cell-types (**Figure 3B, Supplemental Figure S3**), including numerous bleomycin-emergent and/or enriched populations (**Figure 3D-F**). In particular, activated fibroblasts (identified by expression of *Cthrc1* ^21^ were rare in PBS-treated mice, but comprised the majority of fibroblasts recovered from YT^del^ bleomycin treated lungs (**Figure 3C, E**). The increase in activated fibroblast numbers was also accompanied with increased expression of fibrillar collagens *Col1a1*, *Col3a1*, *Col5a3*, *Col6a3* as well as *Fn1* within the activated fibroblast population. There was a shift towards increased *Gdf15+* intermediate alveolar epithelial cells and decreased AT2 cells in the YT^del^ bleomycin treated populations (**Figure 3D,G**). As expected, YT^del^ AT2 cells had reduced Yap/Taz signaling, and after bleomycin exhibited including lower levels of transitional/ intermediate/AT1 cell markers, Yap/Taz binding partner *Nfib*, canonical YT target genes *Ccn1* and *Ccn2*, but increased expression of canonical monocyte chemotactic factor *Ccl2* and Tgfβ-activating integrin αV (*Itgav*) (**Figure 3H,I**). YT^del^ bleomycin injured lungs had increased inflammatory cells overall, with increased monocyte-derived macrophages compared to WT (**Figure 3F, K).** With evidence of exaggerated and persistent inflammation 28 days after bleomycin, we performed flow cytometry to quantify immune subpopulations at the time of peak inflammation (7 days after bleomycin), (**Supplemental Figure S4).** YT^del^ lungs had increased monocytes, neutrophils, and interstitial macrophages 7-days post bleomycin injury (**Figure 3L)**. These findings support the concept that Yap/Taz deletion specifically in AT2 cells leads to abnormal alveolar epithelial cell repair, promotes recruitment of monocyte-derived macrophages, enhances profibrotic activation and proliferation of resident fibroblasts.

**Figure 3:**
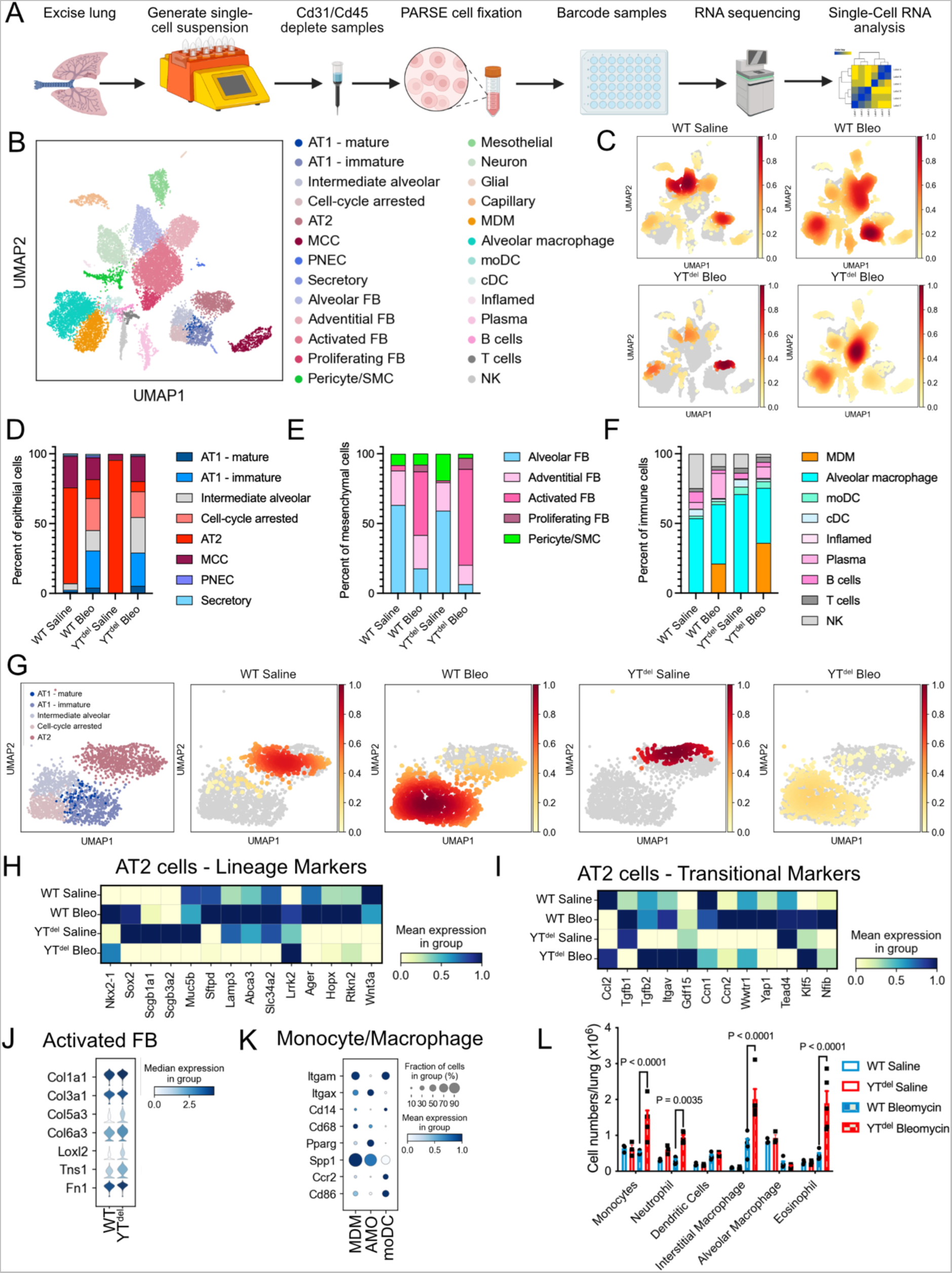
YAP/TAZ deletion prior to bleomycin injury leads to altered epithelial cell populations, activated fibroblast, and disrupted immune response. **A)** Schematic of experimental strategy to generate single-cell RNA analysis from mouse lungs. **B)** UMAP demonstrating clustering of the 26 cell types identified in the mouse lungs. **C)** Density plot showing the representation of each cell type from the respective genotype and treatment groups. **D-F)** Relative proportions of each cell types/states within **D)** epithelial cell populations, **E)** fibroblast populations, and **F)** immune cell populations between each treatment group. **G)** UMAP showing epithelial sub-populations isolated from single-cell suspensions and respective density plots showing relative contribution from each group. **H)** Heat-map showing expression levels of AT2 cell lineage markers in each group. **I)** Heat-map showing transitional cell marker expression in each treatment group. **J)** Violin-plot showing relative expression of activated fibroblast markers in WT or YT^del^ bleomycin treated mice. **K)** Dot-plot showing relative expression of monocyte and macrophage markers in bleomycin treated WT or YT^del^ mice. **L)** Flow Cytometry analysis of immune cell sub-populations present in mice 7 days after bleomycin injury.

### Deletion of Yap/Taz leads to failed alveolar repair and fibrotic remodeling after LPS injury

To determine whether AT2 cell Yap/Taz is more generally required for adaptive alveolar repair, we then performed a series of experiments in which Yap/Taz were deleted 2-weeks prior to intratracheal instillation of 3 mg/kg of *E. coli* lipopolysaccharide (LPS) (**Figure 4A**). By 7-days after injury, while WT mice had largely resolved acute inflammation and injury, YT^del^ mice exhibited accumulation of patchy parenchymal fibrotic remodeling persisted at 28 days post injury, long after LPS injury has typically resolved (**Figure 4B**). Throughout the LPS time-course, YT^del^ mice had more severe and persistent injury in compared to controls, with more damaged alveolar regions at 3, 7, and 14 days that persisted at 28 days post injury (**Figure 4C**). Immunofluorescence analysis of AT1 (Hopx) and AT2 (Sp-C) cell markers demonstrated a persistent loss of AT1 cells at 7 and 28-days post-injury, while there is an initial decrease in AT2 cells at day 7, the AT2 cell population recovers by 28 days post-injury in YT^del^ mice (**Figure 4D,E**). These findings reveal that in the acute LPS injury model, loss of Yap/Taz in AT2 cells leads to a persistent defect in alveolar regeneration that results in increased lung injury, inflammation, and non-resolving alveolar collagen deposition.

**Figure 4:**
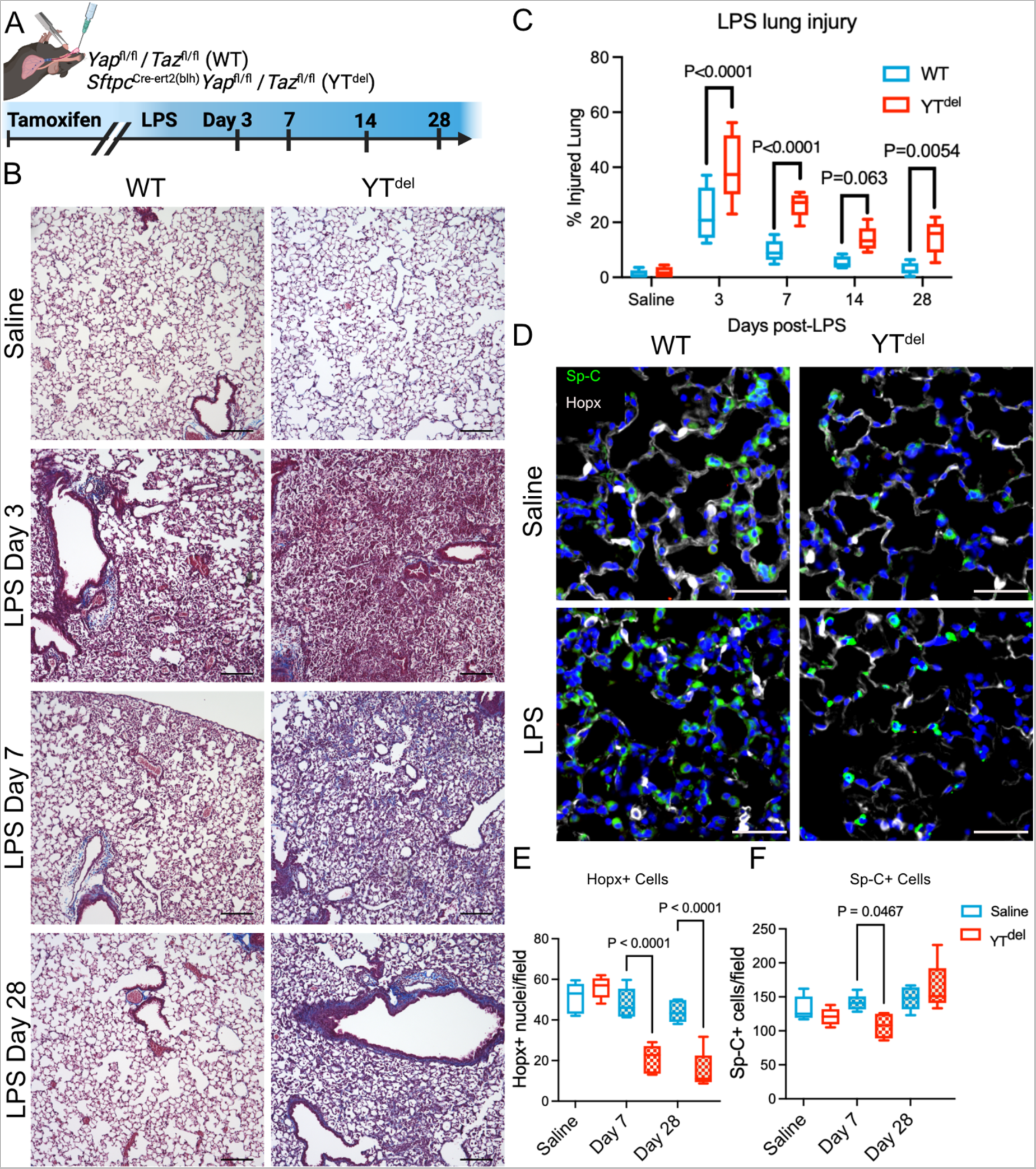
Loss of YAP/TAZ prior to LPS induced lung injury leads to failed alveolar regeneration and increased collagen deposition. **A)** Schematic of LPS lung injury model in which mice were treated with tamoxifen two weeks prior to LPS lung injury. **B)** Masson’s Trichrome staining of wild-type (WT) and YT^del^ mice treated with saline or 3, 7, or 28-days after LPS injury. **C)** Quantification of injured lung area in WT and YT^del^ mice 3 to 28 days after LPS. **D)** Immunofluorescence analysis of Hopx+ (white) AT1 cells and Sp-C+ (green) AT2 cells 7-days after LPS injury. **E, F)** Quantification of Hopx+ and Sp-C+ cells in WT or YT^del^ lungs treated with saline or LPS at 7 or 28-days after injury. Statistics were performed using one-way ANOVA and Tukey’s post-hoc comparison.

### AT2 cell deletion of Yap/Taz disrupts the adaptive innate immune response following LPS

Having observed augmented inflammatory cell recruitment in YT^del^ mice following LPS injury, we then sought to determine the mechanisms through which AT2 cell Yap/Taz regulated acute inflammatory responses. We performed flow cytometry of immune cell populations through a time-course after LPS (**Figure 5A**). At day 3 after LPS, YT^del^ mice had reduced numbers of interstitial macrophages, but increased neutrophils and eosinophils. By 7-days post injury YT^del^ mice had increased interstitial macrophages, neutrophils, and eosinophils indicating prolonged inflammatory response; lymphocyte numbers were similar between groups. These increased inflammatory populations had returned to near WT levels by 14 days after injury (**Figure 5B**, **Supplemental Figure S5**). ELISA analysis of immune cytokines in bronchoalveolar lavage fluid isolated from these mice demonstrated that IL6, CXCL1, and CXCL2 were all increased 3 days post-injury, which were returned to WT levels by day 7 after injury (**Figure 5C**). These data indicated that loss of AT2 cell Yap/Taz worsens alveolar inflammation at least in part via exaggerated expression of neutrophil and macrophage chemotactic factors.

**Figure 5:**
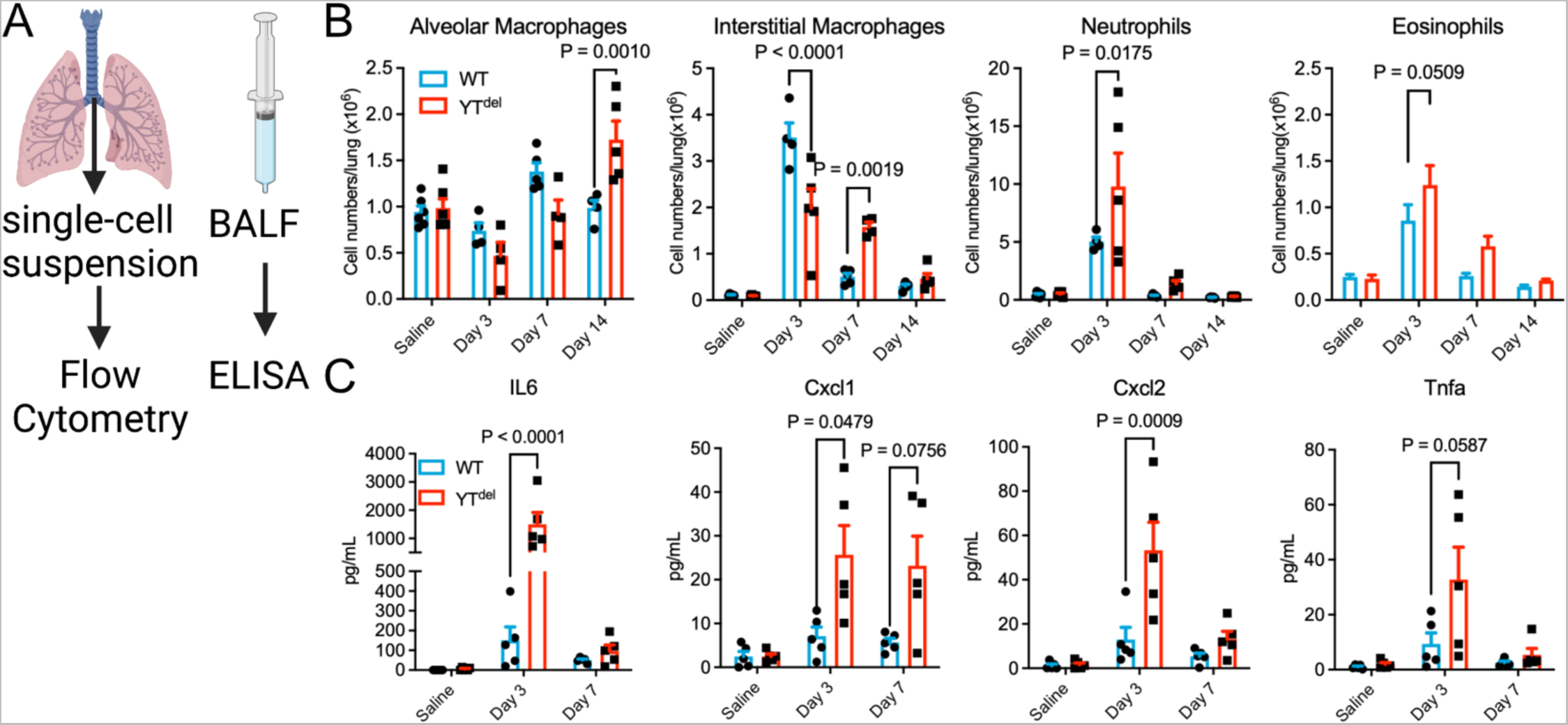
AT2 cell specific deletion of YAP/TAZ leads to altered immune response after LPS injury. **A)** Schematic showing experimental strategy to assess immune response in mice injured with LPS. **B)** Quantification of immune cell types by flow cytometry in saline or LPS treated mouse lungs at 3, 7, or 14-days post injury. **C)** ELISA analysis of immune cytokines isolated from bronchoalveolar lavage fluid from mice 3 or 7 days after LPS injury or saline controls. One-way ANOVA and Tukey’s post-hoc comparison was used to assess statistical significance.

## Discussion

Functional repair and regeneration of the alveolar epithelium is a complex process requiring progenitor cells to proliferate and subsequently mature into structurally and functionally distinct AT2 and AT1 cells. We demonstrated that Yap/Taz activation in AT2 cells is essential for regeneration of both AT2 and AT1 cells during alveolar repair. Further, using complementary models of alveolar injury, we found that AT2 cell Yap/Taz activity restrains innate immune activation, and that without Yap/Taz activation in AT2 cells, ineffective alveolar epithelial repair result in persistent parenchymal injury and fibrosis. Our results here indicate that following injury, YT^del^ lineage-labeled AT2 cells fail to proliferate or give rise to morphologically or transcriptionally mature AT1 cells, suggesting that during injury-repair, at least transient Yap/Taz activity is required to mediate both AT2 proliferative responses and AT1 cell maturation.

Our results build upon and extend recent studies that highlight the complex, dynamic, and cell-type specific roles the Hippo-Yap/Taz pathway plays in lung development, homeostasis and injury-responses. We have previously demonstrated aberrant and persistent YAP activation in the IPF lung epithelium ^6^, while several groups have implicated YAP/TAZ as mediators of fibroblast activation driven by mechanical forces ^22–25^. This has led to interest in YAP/TAZ antagonism as a potential novel antifibrotic strategy in the lung, similar to investigations in other organs ^26–31^. Together with complementary studies from our group and several others, our findings suggest a more nuanced consideration of the Hippo-Yap/Taz pathway is likely required.

During lung development, we and others have previously shown that Yap/Taz is required for AT1 cell commitment ^5, 15, 32, 33^, although recent reports have yielded inconsistent results as to the specific roles of Yap and Taz at different developmental stages and using different models ^4, 34, 35^. Our results here indicate that following single-dose bleomycin or LPS injury, combined deletion of Yap and Taz in AT2 cells prevents AT2 proliferation and AT1 differentiation, indicating that some level of Yap/Taz activity is required for both of these critical aspects of alveolar regeneration. These data complement prior work from our group and others that suggests deletion of Yap/Taz in AT2 cells leads to an “exaggerated AT2 cell phenotype,” ^16^, suggesting that even under homeostatic conditions, low-level Yap/Taz activity may be involved in “priming” AT2 cells for differentiation. In addition, deletion of Yap/Taz in Hopx lineage-labeled cells resulted in reversion to an AT2-like cellular profile ^15^. Together, these findings suggest that Yap/Taz is essential for AT2 cell proliferation, AT1 differentiation, and AT1 cell maturation/maintenance. While it is somewhat counter-intuitive that these distinct processes may all require Yap/Taz, recent studies raise the possibility that different Yap/Taz functions could be regulated through differential interaction partner coupling. This concept is supported by recent work finding dynamic interaction of Yap/Taz with NKX2-1 and/or NFIB and the AT1 associated KLF5 during post-natal lung development ^4, 16, 32, 36, 37^.

We also observed that in two different models of sterile alveolar injury, loss of Yap/Taz activity in AT2 cells was accompanied by increased and persistent alveolar inflammation. These findings are consistent with previous work that found deletion of Yap/Taz leads to increased immune response in *Streptococcus pneumoniae* injured mouse lungs and was associated with suppression of Ikkβ, and upregulation of Nuclear-factor-kappa-beta (Nfkb) signaling ^3^. It is not yet clear whether these impacts on acute inflammation are due to direct effects of Yap/Taz on regulation of inflammatory programs in AT2 cells or are consequences of ineffective alveolar repair. Beyond the role of AT2-cell type specific Yap/Taz activity regulating the immune response of the injured lung, Yap/Taz also plays a role in immune cells through several mechanisms which has recently been reviewed ^38^.

In addition to autonomous effects on epithelial cell fate/function and immune regulation, we also found that deletion of Yap/Taz in AT2 cells worsens fibrotic remodeling following bleomycin injury and was associated with enhanced fibroblast activation and proliferation. More surprisingly, we observed that without AT2 cell Yap/Taz, low-dose LPS led to alveolar collagen deposition and persistent parenchymal remodeling out to at least 28 days after LPS. Together, these data add to a growing body of evidence that suggests specifically that inability to effectively mature AT1 cells is sufficient to promote lung parenchymal fibrosis ^39, 40^, although the specific mechanism through which the pathologic cellular crosstalk drives this process is not yet clear. In the fibrotic lung, there is an increase in stretch response as the lung stiffens during the fibrotic remodeling, and fibroblast-focused studies have shown that Yap/Taz are activated in the presence of mechanical stretch ^22–24, 41^. Additional recent findings have shown that Yap/Taz activation is involved in a stretch associated modulation of chromatin remodeling ^42^ by the extra-cellular matrix, raising the possibility that failure of alveolar epithelial repair alters the local mechanical environment to promote Yap/Taz activation in alveolar fibroblasts and/or adjacent alveolar epithelial cells to promote tissue fibrosis could generate a feed-forward mechanism.

Yap/Taz have also been shown to interact with several other pathways that have been implicated in promoting fibrosis, including Tgfβ ^43, 44^, Wnt ^45–48^, FGF ^49^, Notch ^50^, and mTOR/Pi3K ^51^.

There are several limitations of this data. First, our studies rely on genetic strategies using transgenic mice to modulate Yap/Taz activity in AT2 cells, and the dosing and timing of tamoxifen administration likely impacts the spectrum of recombined cells. Our lineage-tracing studies suggest our recombination efficiency was high in this model, and incomplete recombination would likely bias between-group differences toward the null. There is also inherently limited ability to discern AT2-cell autonomous effects of Yap/Taz deletion from more general consequences of ineffective alveolar repair. We also did not seek to determine separable roles of Yap versus Taz in AT2 cells following injury, as recent reports demonstrate some redundancy in these roles or at least deletion of Yap activates Taz ^34^.

Together, our findings support the concept that the Hippo-Yap/Taz pathway must be tightly regulated to facilitate adaptive repair, and that either failure to activate the pathway to initiate alveolar repair, or failure to down-regulate the pathway to promote alveolar epithelial cell homeostasis both are maladaptive and lead to progressive fibrotic remodeling. These findings collectively support the concept that future therapeutic strategies to modulate YAP/TAZ activity have potential to promote lung regeneration, but that timing and cell targeting considerations will be important.

## Methods

### Animal Husbandry and deletion of YAP/TAZ

*SftpcCre^ert2(blh)^Yap^flox/flox^Taz^flox/flox^* mice were crossed with *Yap^flox/flox^Taz^flox/flox^* mice to generate experimental mouse cohorts similar to our previous post-natal studies ^4^. Cre negative littermates are used for WT controls. *SftpcCre^ert2rosatdTomato^Yap^flox/flox^Taz^flox/flox^* mice were crossed with *Yap^flox/flox^Taz*^flox/flox^ mice to generate mice for lineage tracing studies. *SftpcCre^ert2rosetdTomato^* were used as lineage trace control mice. Mice were administered tamoxifen100mg/kg or corn-oil control by intraperitoneal injection either 2 weeks prior to, or at the same time as respective injury to induce gene deletion and lineage tracing. For assessment of adaptive repair in bleomycin injured mice and as additional control mice for injury experiments, 8-10 week old C57Bl/6J mice were ordered from Jackson laboratory (000664).

### Lung injury

Bleomycin (0.08IU or 0.04IU) or *E. coli* lipopolysaccharide (LPS) (3mg) were suspended in sterile saline and intratracheally administered in 100ul volumes and equivalent volumes of saline were used for controls. Mice were weighed weekly to assess health throughout the experiments. Mice were sacrificed at indicated time-points with the left lung being inflation fixed in 10% buffered formalin and right lung being processed for Sircol collagen analysis or isolation of Cd326+ cells for RNA analysis.

### Immunohistochemical analysis

Lungs were inflation fixed with 10% buffered formalin at 25 cm H20 pressure and fixed at 4 degrees C overnight. Lungs were then processed, paraffin embedded and 5μm sections were used for all histological stains. Immunofluorescence was performed using antibodies (described in **Supplemental table S1**) as follows: Antigen retrieval was done using a rice cooker and 1X Citrate Buffer, then washed in DI water. Slides were blocked in 5% BSA for one hour at room-temperature and then incubated in primary antibodies overnight. Samples were then washed 3X in PBST and then incubated in secondary antibody with DAPI diluted in PBST for 2 hours. Slides were then washed in PBST 3X and cover slips were added with mounting media. For immunofluorescence analysis, samples were imaged in a Keyence BZ-X710 inverted fluorescence microscope, images were taken on a 20X objective of at least 10 non-overlapping fields of view. Image analysis was performed with HALO image analysis software for automated quantification of respective cell states. Histological analysis of lung injury was quantified using ImageJ on samples stained with Masson’s Trichrome stain.

### Single cell RNA sequencing

Single cell suspensions were generated as previously described, in short; Right lung lobes were extracted and incubated in dispase II (Roche), collagenase (Sigma-Aldrich) and DNAse in phenol-free DMEM (Gibco). Lungs were disaggregated using GentleMacs cells dissociator with C-tubes (Miltenyi) and cell suspensions were passed through 100 and 70um filters to isolate single cell suspensions. Once single cell suspensions were achieved, red blood cells were depleted using ACK buffer and cells were fixed following PARSE Biosciences Fixation User Manual V1.3.0 protocol. Libraries were sequenced on a NovaSEQ6000 targeting 50,000 reads per cell, and demultiplexing was performed using the PARSE pipeline (v0.9.6) with default parameters. The read mapping and alignment were based on Mouse GRCm39 genome and GENCODE M28 annotation.

### scRNA-seq analysis

Single-cell RNA-sequencing data were analyzed through a standard Scanpy ^52^ (v1.9.1) workflow similar to our prior work ^53^. Following quality-control filtering (excluding cells with <500 or >5000 genes, and cells with >10% mitochondrial reads), data were normalized and scaled, including mitochondrial and ribosomal percentages as regression variables, followed by principal components analysis, neighborhood graph calculation and UMAP embedding using the first 45 PCs (Principal Components), then recursive leiden-clustering/subclustering. Heterotypic doublet clusters were identified by coexpression of lineage-specific marker genes, and cell annotation was performed manually, informed by published reference datasets ^54, 55^. Cell-type specific differential expression was performed using the wilcoxon test.

### Flow cytometry

To obtain single-cell suspensions from lung tissue, lungs were perfused with sterile PBS, removed *en bloc*, and perfused lungs were digested in RPMI medium containing collagenase XI (0.7 mg/ml; Sigma-Aldrich) and type IV bovine pancreatic DNase (30 µg/ml; Sigma-Aldrich). Red blood cells (RBC) were lysed with RBC Lysis Buffer (BioLegend) as described elsewhere ^56^. Single-cell suspensions were incubated with a Fc receptor block (Cat. 553141, BD Bioscience) to reduce nonspecific anti-body binding. The flow cytometry panel used to identify immune cell subtypes is shown in supplemental figure S3, in short antibodies used in these experiments included CD45-Brilliant Violet 650 (Cat. 103151), Ly6G-APC/Cyanine7 (Cat. 127623), F4/80-PE/Cy5 (Cat. 123111), MerTK-Brilliant Violet 605 (Cat. 151517), CD11c-AF-700 (Cat. 117320), CD11b-PE/Cyanine7 (Cat.101216), MHCII-FITC (Cat. 107605), CD64-APC (Cat. 139306) from Biolegend; SiglecF-PE (Cat. 552126) from BD Bioscience. Dead cells were excluded using DAPI (Cat. MBD0015, Sigma). Flow cytometry was performed using BD LSR II and BD FACS Aria III flow cytometers (BD Bioscience), and data were analyzed with FlowJo software.

### Bronchoalveolar lavage fluid isolation

Intratracheal cannulation was used to wash lungs with 5 times with 1 mL sterile PBS. Immune cell populations were assessed by cytospin and subsequent DiffQuick staining or through the above flow cytometry panel.

### Statistics

Statistics were performed in GraphPad Prism. All bar graph error bars are standard error of the mean (SEM). Whiskers in Box-Whisker plots are min and max values. Statistical significance was determined by one-way ANOVA with Tukey’s post-hoc comparisons. Any deviation in this approach is noted in the respective figure legends. All p-values considered significant (<0.05) are written in the graphs to show comparisons.

### Study Approval

These studies were approved by Vanderbilt University Medical Center’s Institutional Animal Care and Use Committee.

### Data Availability

Raw genomic data are available through the Gene Expression Omnibus (GEO)/Short read archive (SRA) under accession number (pending). Code use for scRNA-sequencing analysis is available at github.com/kropskilab/digiovanni_2023.

### Author Contributions

GTD, WH, TS, TSB, JAK, and JJG designed research studies, GTD, WH, TS, CJT, NMG, UKS, CLC, ASM, MMD, JJ, SSG, and JJG conducted experiments, GTD, WH, CJT, DSN, YL, JAK, and JJG acquired data, GTD, WH, YL, VVP, TSB, JAK, and JJG analyzed data, GTD, TSB, JAK and JJG wrote the manuscript, all authors provided editing to the manuscript.

### Funding

This work was made possible by the Pulmonary Fibrosis Foundation (JJG), by an independent grant from Boehringer Ingelheim Pharmaceuticals, Inc. who provided the financial support. The authors meet criteria for authorship as recommended by the International Committee of Medical Journal Editors (ICMJE) and were fully responsible for all aspects of the study and publication development. This was supported by NIH/NHLBI R01HL153246 (JAK) Vanderbilt Faculty Research Scholars (JJG), P01HL092870 (TSB), T32HL04296 (ASM).

**Supplemental Figure S1: A)** Schematic of bleomycin lung injury, in which mice are treated with tamoxifen 2-weeks prior to injury with a planned 28-day recovery. **B,C)** Survival curves of mice treated with 0.08IU bleomycin (B) or 0.04IU bleomycin (C). Mantel-Cox test was used to determine statistical significance.

**Supplemental Figure S2:** YT^del^ generate fewer and smaller organoids in feeder-free culture. **A.)** Fluorescent image of Tomato lineage labeled AT2 cells cultured in SFFFM media for 14-days. **B.)** Quantification of total number of organoids per well and **C.)** organoid size analysis of organoids larger than 25μm. Statistical analysis performed with an unpaired t-test. Scale bar represents 25μm.

**Supplemental Figure S3:** Cell types identified in single-cell RNA sequencing analysis and expression of cell-type specific markers.

**Supplemental Figure S4**: Flow Cytometry gating strategy of immune cells isolated from mouse lung single-cell suspensions.

**Supplemental Figure S5:** Flow cytometry of total numbers of monocytes and dendritic cells isolated from LPS injured lungs at day 3, 7, and 14 after injury. Cell numbers were not significantly different based on genotype determined by one-way ANOVA and Tukey’s post-hoc comparison.

**Supplemental Table S1:** Antibodies used for immunofluorescence analysis with respective catalog information and concentration used.

