## Supplemental Figures for "Epithelial Yap/Taz are required for functional alveolar regeneration following acute lung injury"

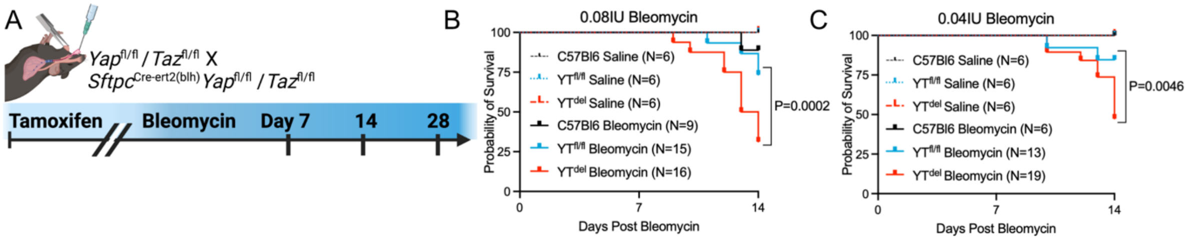


**Supplemental Figure S1: A)** Schematic of bleomycin lung injury, in which mice are treated with tamoxifen 2-weeks prior to injury with a planned 28-day recovery. **B,C)** Survival curves of mice treated with 0.08IU bleomycin (B) or 0.04IU bleomycin (C). Mantel-Cox test was used to determine statistical significance.

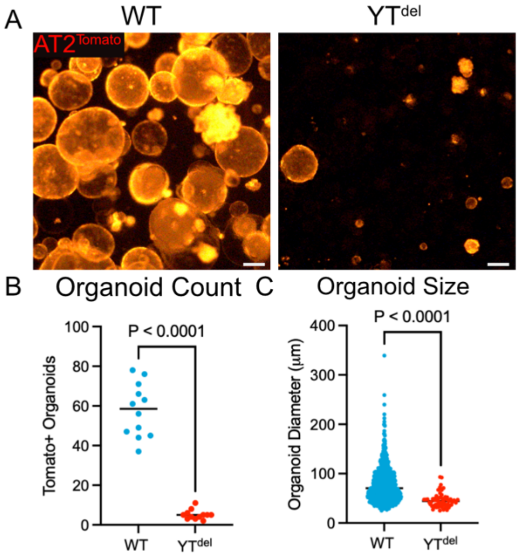


**Supplemental Figure S2:** YT$^{del}$ generate fewer and smaller organoids in feeder-free culture. **A.)** Fluorescent image of Tomato lineage labeled AT2 cells cultured in SFFFM media for 14-days. **B.)** Quantification of total number of organoids per well and **C.)** organoid size analysis of organoids larger than 25μm. Statistical analysis performed with an unpaired t-test. Scale bar represents 25μm.

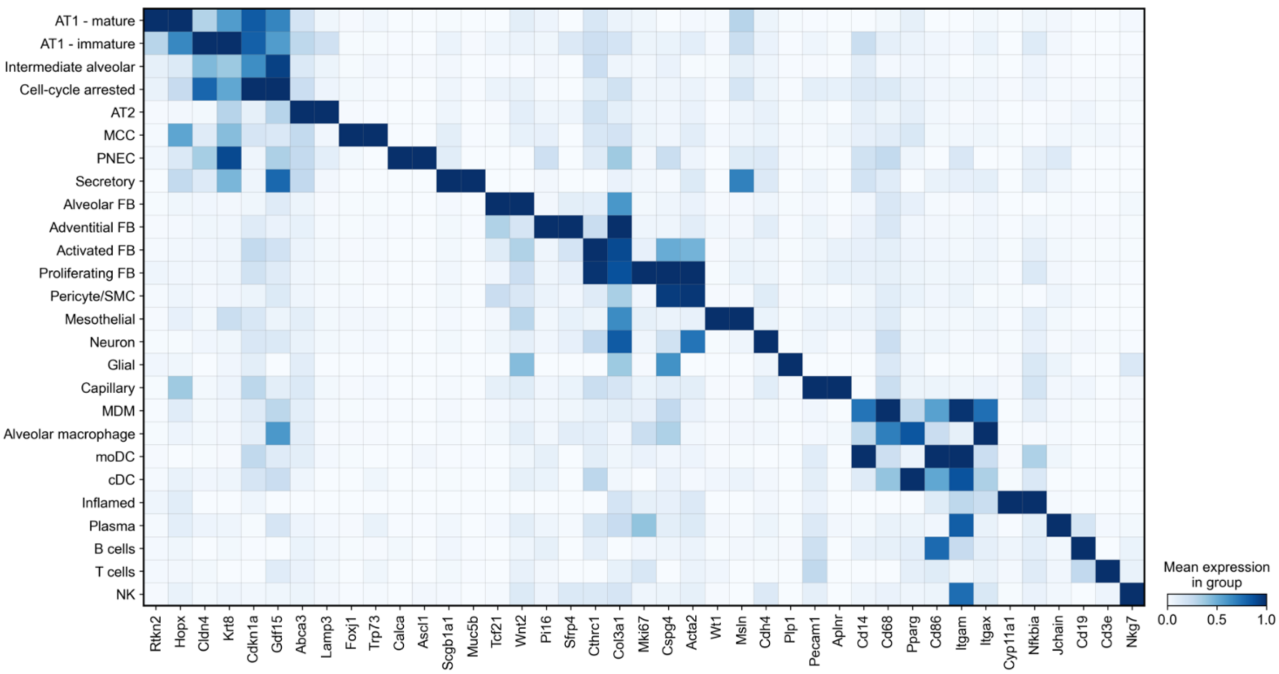


**Supplemental Figure S3:** Cell types identified in single-cell RNA sequencing analysis and expression of cell-type specific markers.

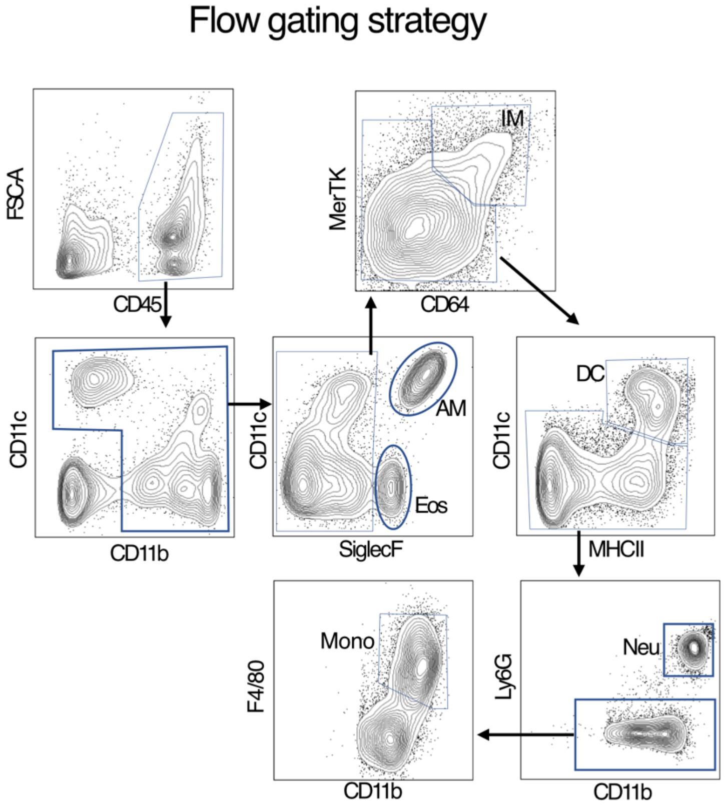


**Supplemental Figure S4**: Flow Cytometry gating strategy of immune cells isolated from mouse lung single-cell suspensions.

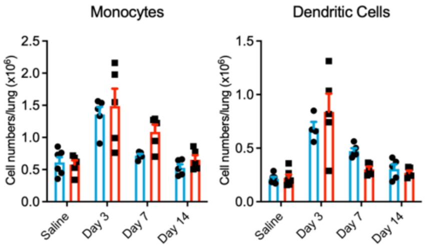


**Supplemental Figure S5:** Flow cytometry of total numbers of monocytes and dendritic cells isolated from LPS injured lungs at day 3, 7, and 14 after injury. Cell numbers were not significantly different based on genotype determined by one-way ANOVA and Tukey's post-hoc comparison.

| Antibody | Company | Host Species | Catalog # | Concentration |
|---|---|---|---|---|
| SPC | Seven Hills Bioreagents<br><br>Abcam | Rabbit<br><br>Rabbit | WRAB-9337<br><br>AB90716 | 1:400<br><br>1:100 |
| YAP | Seven Hills Bioreagents<br><br>Cell Signaling Technology | Rabbit<br><br>Rabbit | WRAB-1549<br><br>4912S<br>14074S | 1:100<br><br>1:100<br>1:100 |
| TAZ | Cell Signaling Technology | Rabbit | 83669S | 1:100 |
| HOPX | Santa Cruz Biotechnology | AF647 Conj. | sc-398703 | 1:100 |
| SCGB1A1 | Santa Cruz Biotechnology | AF488 Conj. | sc-365992 | 1:500 |
| Ki67 | Cell Signaling Technology | AF647 Conj. | 12075S | 1:100 |
| TdTomato | SICGEN | Goat | AB8181-200 | 1:500 |
| AGER | R&D Systems | Rat | MAB1179 | 1:200 |

**Supplemental Table S1:** Antibodies used for immunofluorescence analysis with respective catalog information and concentration used.
